# Speech motor cortex enables BCI cursor control and click

**DOI:** 10.1101/2024.11.12.623096

**Authors:** Tyler Singer-Clark, Xianda Hou, Nicholas S. Card, Maitreyee Wairagkar, Carrina Iacobacci, Hamza Peracha, Leigh R. Hochberg, Sergey D. Stavisky, David M. Brandman

## Abstract

Decoding neural activity from ventral (speech) motor cortex is known to enable high-performance speech brain-computer interface (BCI) control. It was previously unknown whether this brain area could also enable computer control via neural cursor and click, as is typically associated with dorsal (arm and hand) motor cortex. We recruited a clinical trial participant with ALS and implanted intracortical microelectrode arrays in ventral precentral gyrus (vPCG), which the participant used to operate a speech BCI in a prior study. We developed a cursor BCI driven by the participant’s vPCG neural activity, and evaluated performance on a series of target selection tasks. The reported vPCG cursor BCI enabled rapidly-calibrating (40 seconds), accurate (2.90 bits per second) cursor control and click. The participant also used the BCI to control his own personal computer independently. These results suggest that placing electrodes in vPCG to optimize for speech decoding may also be a viable strategy for building a multi-modal BCI which enables both speech-based communication and computer control via cursor and click.

## 1. Introduction

The pathway from the brain to the muscles can be disrupted by neurological disease or injury such as stroke or amyotrophic lateral sclerosis (ALS), resulting in an inability to move or communicate. Intracortical brain-computer interfaces (BCIs) are surgically-implanted devices which bypass the damaged pathway by reading neural signals directly from the brain and producing an output on the user’s behalf. For the past few decades, a major focus of the BCI field has been decoding neural activity associated with attempted movements to control a computer cursor.^1–17^ By controlling a computer cursor with their neural signals, a person with paralysis can type sentences using an on-screen keyboard, send emails and text messages, or use a web browser and many other software applications.^8,12,14^ Most BCIs of this class have been driven by neural activity in *dorsal* motor cortical areas such as the “hand knob” area of dorsal precentral gyrus (dPCG) and are controlled by the user attempting or imagining *arm or hand* movements.^5,6,8–12^ In recent years, speech BCIs have emerged as a viable path toward restoring fast, naturalistic communication for people with paralysis by instead decoding attempted *speech* movements.^18–23^ In contrast to hand-based BCIs, speech BCIs have typically been driven by neural activity in sensorimotor cortical areas further *ventral* such as middle precentral gyrus (midPCG) and ventral precentral gyrus (vPCG) which are most often associated with production of orofacial movements and speech.^18,20–22^ Speech BCIs far outperform cursor BCIs with regard to communication rate,^10,20^ but are not as well-suited for general-purpose computer control. Though it is possible to implant electrodes in both dorsal and ventral areas, in some patients it may be impractical or infeasible. At first glance, this appears to lead to a difficult decision for prospective neuroprosthesis users and developers: should the surgeon implant intracortical electrodes into dPCG to optimize for computer control, or into vPCG to optimize for speech decoding?

However, motor cortical neural tuning along the precentral gyrus is not as body-part-specific as was once believed. dPCG exhibits modulation and some specificity to speech and other non-hand motor imageries,^24–27^ and vPCG exhibits modulation and some specificity to hand and other non-speech motor imageries.^27^ This begs the question whether vPCG contains enough hand-related (or potentially effector-agnostic^26^) movement intention information to drive a cursor BCI. This would make it a desirable implant target for a multi-modal BCI (a speech BCI *and* a cursor BCI) that maximizes speech-based communication accuracy while still providing cursor-based computer control.

We explored the possibility that vPCG supports such a multi-modal BCI with a BrainGate2 clinical trial participant who had four intracortical microelectrode arrays implanted in vPCG. A previous study demonstrated that decoding this participant’s neural signals from these arrays enabled a highly accurate speech BCI.^22^ Here, we engaged the participant in BCI cursor control tasks and trained decoders to map his neural activity patterns to cursor velocity and click. Similar to cursor BCIs driven by dPCG, our cursor BCI driven by vPCG proved to be rapidly-calibrating (the participant acquired his first target 40 seconds into his first-ever usage of a cursor BCI) and achieved a peak performance comparable to previous intracortical BCI cursor control studies (up to 3.16 bits per second, average 2.90). We found that most of the information driving cursor control came from one of the participant’s four arrays, and click information was available on all four arrays. Anticipating a multi-function use case, we also measured the impact of the participant speaking while simultaneously controlling the cursor. We found that, in contrast to dPCG cursor BCIs,^28^ speaking did transiently perturb the cursor. Finally, the BCI enabled the participant to neurally control his personal desktop computer to independently perform his own desired consumer application activities.

## 2. Methods

### 2.1. Participant and approvals

This research was conducted as part of the BrainGate2 clinical trial (NCT00912041). Permission for the trial was granted by the U.S. Food and Drug Administration under an Investigational Device Exemption (Caution: Investigational device. Limited by federal law to investigational use). This study was approved by the Institutional Review Board at Mass General Brigham (#2009P000505) and at the University of California, Davis (protocol #1843264). This manuscript does not report any clinical trial-related outcomes; instead, it describes scientific and engineering discoveries that were made using data collected in the context of the ongoing clinical trial.

This report includes data from one participant (‘T15’), a 45-year old left-handed man with ALS. At the time of data collection, T15 was tetraplegic and had severe dysarthria. He retained eye movement, impaired neck movement, and limited orofacial muscle control. T15 gave informed consent for this research to be conducted and published, including video. Data reported in this study were collected between day 39 and day 468 post-implant. All research sessions were performed at the participant’s home.

### 2.2. Neural recordings

We surgically implanted four 64-electrode Utah arrays (NeuroPort Array, Blackrock Neurotech, Salt Lake City, Utah) into T15’s vPCG. Precise target locations were informed by preoperative MRI scans and alignment of T15’s brain with the Human Connectome Project^29^ cortical parcellation (Fig. 1b). See Card et al. 2024 Supplementary Appendix S1.02 for further detail.^22^ The raw voltage signals from the 256 electrodes were collected at 30 kHz and preprocessed by a neural data acquisition system (NeuroPort System, Blackrock NeuroTech). Each signal was then bandpass filtered between 250 and 5000 Hz. We applied Linear Regression Referencing (LRR) to each array’s 64 signals to filter out common noise seen on multiple electrodes.^30^

**Figure 1.**
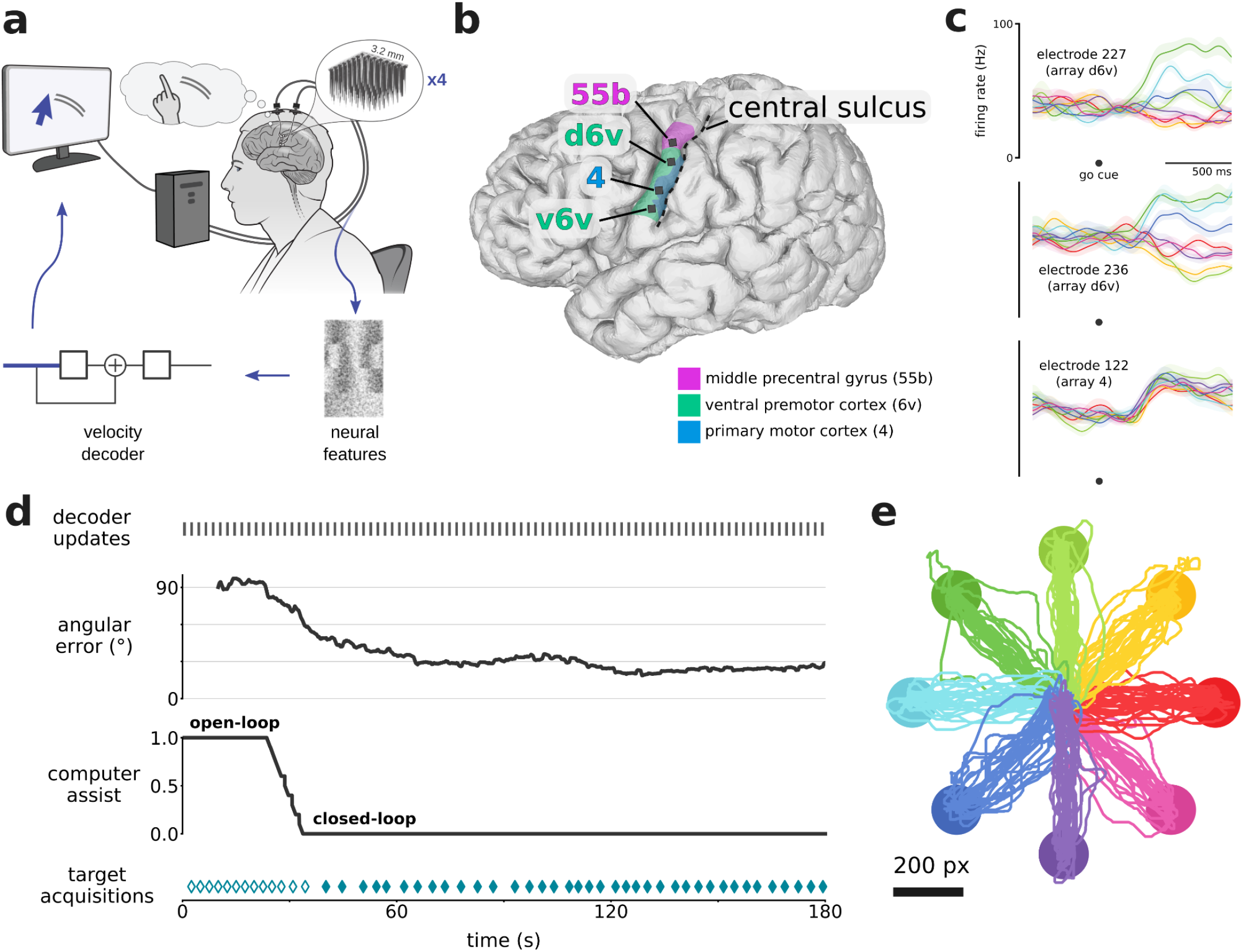
Rapid calibration of cursor control during first-ever usage. **a**. Schematic of the cursor BCI. Neural activity was recorded by four 64-electrode arrays located in the left ventral precentral gyrus. Neural features were calculated in bins of 10 ms. A linear velocity decoder predicted the participant’s intended velocity and moved a computer cursor accordingly. **b**. 3-D reconstruction of T15’s brain (left view), with implanted Utah array locations (black squares) and brain areas (magenta, green, blue regions) overlaid. Brain areas were estimated through alignment of T15’s brain with the Human Connectome Project cortical atlas via preoperative MRI scans. **c**. Trial-averaged firing rates (mean ± s.e.) recorded from three example electrodes during the Radial8 Calibration Task. Activity is aligned to when the target appeared (which was also the go cue), and colored by cued target direction (same colors as in panel e). Multiple arrays showed cursor-related modulation, and some electrodes showed directional tuning. Firing rates were Gaussian smoothed (std 50 ms) before trial-averaging. **d**. Timeline of the first 3 minutes that T15 ever used a cursor BCI. Gray marks every few seconds (top) indicate updates to the cursor decoder. Angular error between the vector toward the target and the vector predicted by the decoder is plotted as a rolling average (20 seconds). The task began in open-loop (assist = 1.0) and gradually transitioned to closed-loop (assist = 0.0) as angular error lowered. Teal diamonds (bottom) indicate successful target acquisitions (reaching the target and dwelling for 0.8 seconds). Filled diamonds indicate fully closed-loop trials. **e**. Center-out-and-back cursor trajectories (224 trials) during T15’s first six Radial8 Calibration Task blocks, colored by cued target direction.

Every 1 ms, two neural features (threshold crossings and spike band power) were calculated per electrode, yielding 512 neural features per ms. Each electrode’s threshold was defined as -4.5 times that electrode’s root mean square (RMS) voltage value, and a threshold crossing was registered in a given 1 ms window if the signal reached that electrode’s threshold. Spike band power was the square of the bandpass filtered signal, averaged in each 1 ms window. Each of the 512 neural features was binned into 10 ms bins (threshold crossings were summed, spike band power was averaged). This time series of 512-D binned neural features vectors was the input for all online control and decoding steps presented in this study (referred to as *f*_*t*_ in Sections 2.4. and 2.5.).

Decoders used both threshold crossings and spike band power for all closed-loop experiments in this study, but post-hoc analyses indicate that comparable performance could have been achieved using only spike band power (Supp. Fig. 1).

### 2.3 Behavioral tasks

We engaged T15 in three formal cursor BCI tasks: a ‘Radial8 Calibration Task’ (Supp. Movie 1), a ‘Grid Evaluation Task’ (Supp. Movie 2), and a ‘Simultaneous Speech and Cursor Task’ (Supp. Movie 3). T15 also engaged in self-directed use of the BCI to control his personal computer. Results reported in this study are based on data from six data collection sessions: one ‘First-ever Usage Session’, four ‘Evaluation Sessions’, and one ‘Simultaneous Speech and Cursor Session’. The sessions in which each task was conducted are listed in Table 1.

**Table 1.** Session days, with instructed imageries and decoder parameters. β are *r* are in speed units of screen height / second. α and *p* are unitless. β is specified for sessions where we normalized the tuning matrix such that β was in meaningful units. *r* and *p* are specified for sessions where the nonlinear speed adjustment was used (i.e., *p* ≠ 1. 0).

| Post-implant day | Session type | Task | Instructed cursor imagery | Instructed click imagery | Cursor decoder parameters |
| --- | --- | --- | --- | --- | --- |
| 39 | First-ever Usage | Radial8 Calibration | Right Hand Mouse, Left Hand Mouse, Right Hand Paintbrush, Left Hand Paintbrush, Tongue Sticking Out, Tongue Inside Mouth | n/a | $\alpha = 0.9$ |
| 171 | Evaluation | Radial8 Calibration | Head Mouse / Intuition | Close Mouth, Close Right Hand | $\alpha = 0.9$<br>$\beta = 0.2, 0.15$ |
|  |  | Grid Evaluation | " | " | " |
| 202 | Simultaneous Speech and Cursor | Radial8 Calibration | Right Hand Mouse | n/a | $\alpha = 0.9$<br>$\beta = 0.2$ |
| | | Simultaneous Speech and Cursor | " | " | $\alpha = 0.9$<br>$\beta = 0.15$ |
| 233 | Evaluation | Radial8 Calibration | Right Hand Mouse | Close Right Hand | $\alpha = 0.93$<br>$\beta = 0.16$ |
| | | Grid Evaluation | " | " | $\alpha = 0.9$<br>$\beta = 0.2$ |
| 339 | Evaluation | Radial8 Calibration | Right Hand Mouse | Close Right Hand | $\alpha = 0.93$<br>$\beta = 0.22$<br>$r = 0.4$<br>$p = 1.4$ |
|  |  | Grid Evaluation | " | " | " |
| 468 | Evaluation | Radial8 Calibration | Intuition | Intuition | $\alpha = 0.92$<br>$\beta = 0.13$<br>$r = 0.05$<br>$p = 1.5$ |
|  |  | Grid Evaluation | " | " | " |

#### 2.3.1. Radial8 Calibration Task

The Radial8 Calibration Task was used to train decoders in all sessions. The cursor workspace was presented on a computer monitor in front of the participant. The basic structure was similar to that of many previous cursor BCI studies.^8,11,31^ The workspace contained eight circular gray targets arranged around a circle in 45° intervals, a ninth target in the center, and a circular white cursor which could move. Each trial began with one target turning green (the ‘go cue’). The participant was instructed to attempt a movement (the ‘instructed cursor imagery’) as if he were moving the cursor toward the cued target. For blocks where click was involved, the participant was instructed to attempt another movement (the ‘instructed click imagery’) when the cursor was touching the target. The trial ended when the participant selected the cued target with the cursor, or when a 10-second limit was reached. The cued target was selected when the cursor dwelled on it (during First-ever Usage Session, and Simultaneous Speech and Cursor Session) or clicked on it (during Evaluation Sessions). The Radial8 Calibration Task’s dwell requirement was 0.8 seconds (during First-ever Usage Session) or 1.0 seconds (during Simultaneous Speech and Cursor Session). The order of cued targets alternated between the center target and an outer target (‘center-out-and-back’). For the outer targets, the order was pseudorandom in that each sequence of eight was a random permutation of all of them. There was no intertrial period or delay period before the go cue (i.e., the next target appeared as soon as the previous trial ended).

T15 performed the task in 3-minute blocks, each of which may contain multiple phases of how much assistance was provided to the participant (Fig. 1d), as described below. To make the user’s decoder calibration experience smoother, we developed a procedure to gradually provide more control to the user based on performance evaluations of a cursor decoder (model and training described in Section 2.4.) running in the background in real time. When beginning a series of calibration blocks, the first seconds of the first block was an ‘open-loop’ phase, in which the cursor automatically moved to each cued target under full computer assist (velocity assist = 1.0), i.e., no neural control. In parallel, every 2-3 seconds, a cursor decoder was trained to map neural data to intended cursor velocity, using all the training data in the block up to that point. Even though the decoder was not being used to control the cursor during this phase, the decoder was still applied to incoming 10 ms bins of neural data to yield predicted velocity vectors. Using these predictions, every 1 second, in the background we computed the angular error (the angle between each predicted velocity vector and the vector from cursor to target, where chance is 90°) over a 20 second sliding window. If the angular error was below 80°, computer assist would begin to lower and the cursor’s movement each frame would be a weighted sum of the decoder’s predicted velocity vector and a vector directly toward the target. If angular error lowered to 60°, cursor movement would be controlled only by the user, with no computer assist (velocity assist = 0.0), representing ‘closed-loop’ cursor control. The technique of frequently updating decoder parameters online follows a previously-demonstrated rapid calibration protocol.^11,32^

We also developed an independent and parallel procedure for calibration of a ‘click’ decoder. Click assist was binary (either on or off). With ‘click assist’ on, the computer automatically registered a click once the cursor was on the target for 1.5 seconds. The click decoder was evaluated according to on-target clicks and off-target clicks that would have occurred had the decoder been used. In an 8-trial sliding window, if 7 or more targets would have been clicked and the number of off-target clicks would have been 4 or fewer, click assist was turned off.

The instructed cursor imageries and instructed click imageries used in each session are listed in Table 1.

#### 2.3.2. Grid Evaluation Task

To quantify the participant’s cursor and click performance using the BCI, we used the Grid Evaluation Task. This task followed the standardized task paradigm employed by previous cursor BCI studies^3,5,10,33,34^ to yield a bitrate measured in bits per second (bps). The cursor workspace contained a grid (either 6×6 or 14×14) of square gray targets, and a circular white cursor which could move. As in the Radial8 Calibration Task, each trial began with a random target turning green, and the participant was instructed to use the instructed cursor imagery and instructed click imagery to move to and select the cued target. The trial ended when the participant selected any target (correct or incorrect), or when a 10-second limit was reached. Selecting the wrong target or timing out were both considered a failed trial and immediately triggered a failure sound. Selecting the correct target was considered a successful trial and immediately triggered a success sound. The target on which the cursor began was disabled from being selected for 2.0 seconds or until the cursor left the target. A thin area (43 px wide) surrounding the grid and a gap (1 px wide) between grid cells could be clicked without triggering a selection.

T15 performed the task in 3-minute blocks. The cursor was under closed-loop neural control the entire block (no velocity or click assist), using a fixed cursor decoder and click decoder which were trained earlier in the session using the Radial8 Calibration Task.

The instructed cursor imageries and instructed click imageries used in each session are listed in Table 1.

#### 2.3.3. Simultaneous Speech and Cursor Task

To measure the impact of the participant speaking while controlling the cursor, we used the Simultaneous Speech and Cursor Task: a variation of the Radial8 Calibration Task that included speech prompts (Fig. 4a). Targets were arranged the same as in the Radial8 Calibration Task. However, each trial began with one target turning red (‘target presentation’) and a word simultaneously appearing above the cursor’s starting location (‘word presentation’). The prompted word was also played aloud using text-to-speech. After a randomized 1.3-1.7 second ‘delay period’, the cued target turned from red to green (‘cursor go cue’) and the prompted word disappeared, initiating the ‘move period’. The participant used the instructed cursor imagery (‘Right Hand Mouse’) to move the cursor to the target and dwell on it to select it. Between trials was an intertrial period of 1.0 seconds.

There were two trial types in every block: ‘yes beep’ trials containing an auditory beep (the ‘speech go cue’) and ‘no beep’ trials containing no beep. In ‘yes beep’ trials, the beep occurred at a random time during the move period or dwell period (either 0.3-2.0 seconds after the cursor go cue, or 0.1 seconds after first entering the target). The dwell requirement was long (1.5 seconds) so that, even for speech go cues that occurred in the dwell period, the participant had time to react and speak before the trial ended.

To further test whether performance was affected by having to attend (or not) to the speech go cue, there were also two block types: in ‘speak if beep’ blocks the participant was instructed to speak the current trial’s prompted word when they heard the beep, whereas in ‘never speak’ blocks the participant was instructed not to speak even if there was a beep (i.e., the participant could ignore all beeps during this block type). With two block types and two trial types, there were four total trial conditions: ‘speak if beep - yes beep’, ‘speak if beep - no beep’, ‘never speak - yes beep’, and ‘never speak - no beep’. As described, only the ‘speak if beep - yes beep’ condition required speaking. This task design was similar to that of a previous study that examined interference between speaking and cursor control using electrodes in dPCG.^28^

### 2.4. Neural cursor decoder

#### 2.4.1. Neural feature preprocessing

Each neural feature in *f*_*t*_ was z-scored according to the data used for training the current decoder, and clipped at ±5. Electrodes with consistently low activity during calibration (mean firing rate <5 Hz) were excluded from training and inference. During inference, for bins where noise was detected on an electrode (4 out of the last 5 bins with firing rate ≥500 Hz, or 4 out of the last 5 bins with spike band power ≥25000 μV^2^), that electrode’s features were set to their mean values.

#### 2.4.2. Cursor decoder model

The cursor decoder was a linear velocity decoder with temporal smoothing,^35^ summarized by the equation

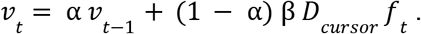

Briefly, *f*_*t*_ is the input (512-D neural features vector) and *v* is the output (2-D cursor velocity vector). This equation defines cursor velocity as a weighted sum of the previous time bin’s velocity and a new velocity predicted from the neural data. The balance between these two terms is set by α, which may be set within the range [0, 1). The left term (α *v* _*t*−1_) provides temporal smoothing. The higher the value of α, the smoother the cursor’s trajectory, but the longer it will take to switch direction. The parameter β is a speed gain, which may be set within the range (0, ∞). The higher the value of β, the faster the cursor will move, but this can make overshooting the target more likely.

*D*_*cursor*_ is the ‘tuning matrix’ consisting of each neural feature’s (scaled) preferred direction,^36^ computed during the training procedure described in Section 2.4.3 (i.e., *D*_*cursor*_ is updated every few seconds during the Radial8 Calibration Task).

One final step was applied to *v*_*t*_ before moving the cursor: a nonlinear speed adjustment,^13^ which made slow velocities slower, and fast velocities faster. The actual cursor velocity output was

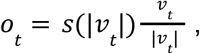

*w*here *s* is a scalar function, so *o*_*t*_ is the same direction as *v*_*t*_ but its speed is adjusted based on the decoded speed itself |*v*_*t*_ |. Specifically,

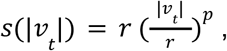

*w*here the parameter *r* is the inflection point between speeds that are slowed down and speeds that are sped up, and the parameter *p* is the degree to which speeds are slowed down or sped up. *p* is configurable within the range [1, ∞), where 1 means no adjustment and higher means more substantial adjustment. The motivation for using this nonlinear speed adjustment was to improve the BCI user’s ability to stop and select small targets as well as their ability to traverse far distances quickly. The parameter *r* was not explicitly parameterized in the original description of the speed adjustment formula,^13^ where it was implicitly hard-coded to *r* = 1.

The specific values used for α, β, *r*, and *p* in each session are listed in Table 1.

#### 2.4.3. Cursor decoder training

We used linear regression to calculate the tuning matrix *D*_*cursor*_, which maps each neural features vector *f*_*t*_ into 2-D cursor velocity space. The inputs were individual time bins (10 ms) of neural features and the outputs were the {*x, y*} components of the participant’s intended velocity at each time bin. We used unit vector intention estimation,^37^ which assumes intended speed is constant while the cursor is far from the target and excludes time bins where the cursor is near the target. An exception to this is that during the First-ever Usage Session, we used position error intention estimation,^37^ which assumes intended speed scales with the cursor’s distance from the target. The training data included samples from all phases (open-loop, transition, and closed-loop^38–40^) of the Radial8 Calibration Task.

The linear regression was performed every 2-3 seconds and incorporated all the training data in the current consecutive series of Radial8 Calibration Task blocks that used a consistent instructed cursor imagery. The active cursor decoder was immediately updated with the new tuning matrix *D*_*cursor*_ each time the training was performed (i.e., during calibration blocks, the decoder weights were updated many times throughout the block, even during ongoing cursor movements).

### 2.5. Neural click decoder

#### 2.5.1. Click decoder model

The click decoder was a linear classifier with temporal smoothing. Each time bin of neural features *f*_*t*_ was multiplied by a tuning matrix *D*_*click*’_ to yield a vector of logits for the candidate classes (‘no click’ or ‘click’). Normalizing this vector so it sums to 1 yields probabilities for the two candidate classes. Each time bin, the predicted class was chosen as the class with the largest probability. Smoothing was implemented by the decoder refraining from actually outputting a click until a portion *T* of predicted classes within a time window of width *L* were ‘click’. This helped reduce errant clicks due to decoding noise. We also employed a cooldown window of width *C* after a click was performed during which another click could not be output.

For all sessions, we set *T* = 0. 8, *L* = 70 ms, and *C* = 1. 0 seconds.

#### 2.5.2. Click decoder training

We used logistic regression to calculate the tuning matrix *D*_*click*_, which maps each neural features vector *f*_*t*_ to 2-D class logit vectors. The inputs were individual time bins (10 ms) of neural features and the output labels were ‘no click’ or ‘click’. Time bins labeled ‘click’ came from the window 0.5-1.0 seconds after the cursor entered the cued target, each time the cursor remained on the target for at least 0.5 seconds. For all other time bins, if the cursor was not on the cued target they were labeled ‘no click’, and if the cursor was on the cued target they were ignored.

Similar to the velocity decoder, the click decoder logistic regression was performed every 2-3 seconds and incorporated all the training data in the current consecutive series of Radial8 Calibration Task blocks that used a consistent instructed click imagery. The active click decoder was immediately updated with the new tuning matrix *D*_*click*_ each time the training was performed.

### 2.6. Software pipeline for experiments

Feature extraction, behavioral tasks, and neural decoders were implemented in Python 3.8 and 3.9, and were run using the BRAND framework.^41^ BRAND encourages modularizing the pipeline into reusable components (processes) called BRAND nodes which use Redis for inter-process communication and logging. Calculation of the decoders’ tuning matrices used Python package `scikit-learn` (1.1.1).^42^

Feature extraction, behavioral tasks, and neural decoders ran on commercially-available computing hardware residing on a wheeled cart near the participant. To move and click the native macOS cursor on T15’s personal Mac computer, we ran a lightweight application on T15’s computer itself. It received the outputs of the BCI’s cursor decoder and click decoder, and then moved and clicked the native cursor using Python package `pynput`.

### 2.7. Data and code availability

Neural features, cursor task data, and code will be available upon publication.

## 3. Results

### 3.1. Rapid calibration during first-ever cursor BCI usage

We first measured speed of onboarding, i.e., how long it took a *first-time* cursor BCI user to gain neural control of a cursor.^11^ During T15’s first-ever cursor BCI task (Fig. 1d), he spent 25 seconds in the open-loop phase (velocity assist = 1.0). At 25 seconds, the decoder’s average angular error (using a 20 second sliding window) reached 80°, at which point velocity assist began to lower as angular error lowered. Between 25 and 34 seconds, angular error dropped to 60° and velocity assist lowered to 0.0, beginning the fully closed-loop phase. At 40 seconds, T15 acquired his first target fully using neural control.

### 3.2. Cursor and click performance

In later research sessions, we supplemented the continuous cursor control with a discrete click. All four arrays contained electrodes which modulated to attempted click (Fig. 2a). We evaluated the BCI’s cursor and click performance in terms of bitrate using the Grid Evaluation Task (Fig. 2b, Supp. Movie 2). Blocks 1-8 used a 6×6 grid, whereas blocks 9-17 used a 14×14 grid and lowered *r* to 0. 05 in order to better leverage the nonlinear speed adjustment. These changes made a substantial difference so our assessments segregate the Grid Task Evaluation blocks into “preliminary” blocks (1-8) and “optimized” blocks (9-17). T15’s average bitrate for the optimized blocks was 2.90 ± 0.16 bps (mean ± s.d.) and for the preliminary blocks was 1.67 ± 0.21 bps (Fig. 2e). T15’s single-block maximum was 3.16 bps.

**Figure 2.**
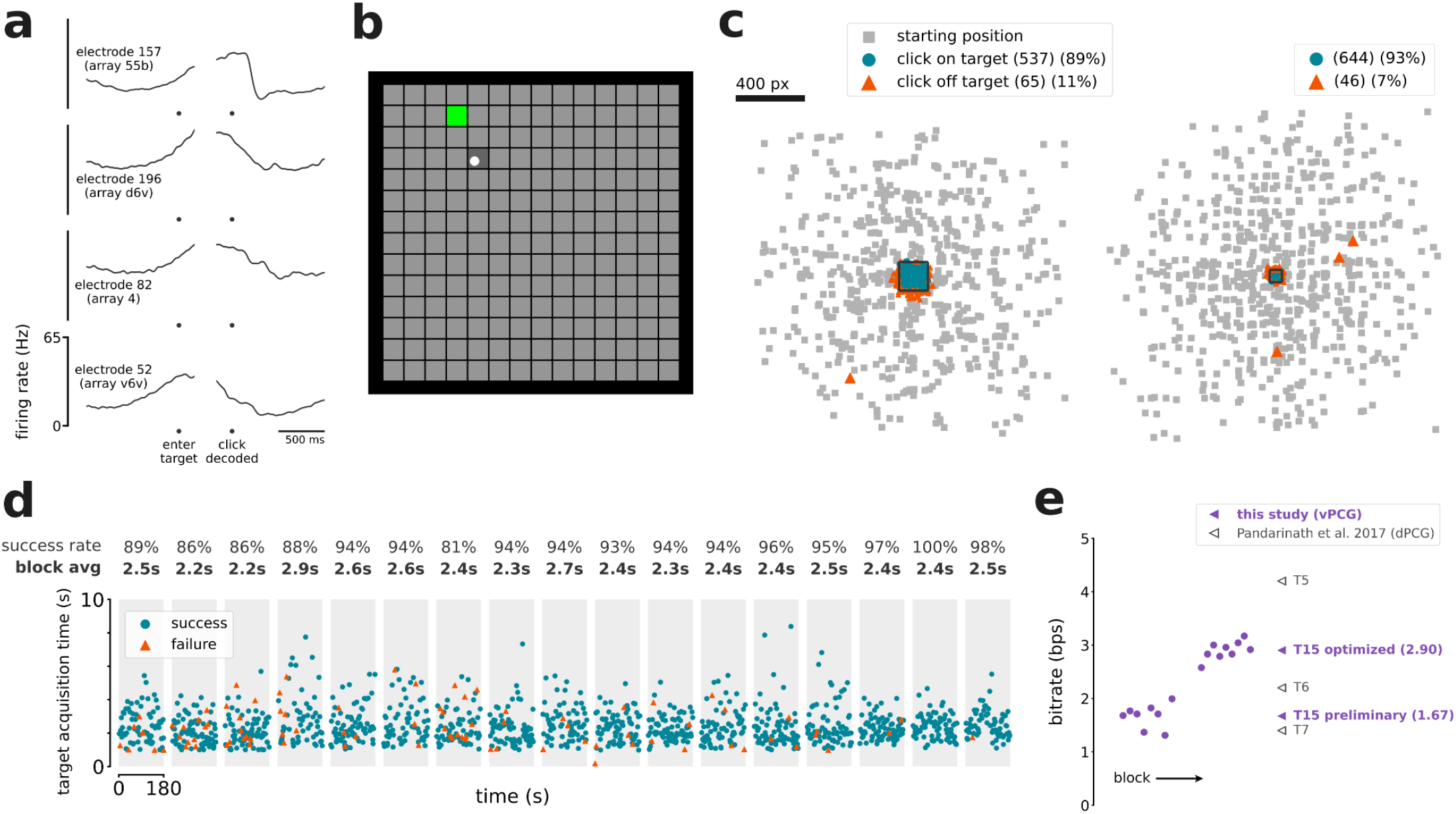
Accurate cursor control and click. **a**. Trial-averaged firing rates (mean ± s.e.) recorded from four example electrodes during the Grid Evaluation Task. Activity is aligned to when the cursor entered the cued target (left), and then to when the click decoder registered a click (right). Firing rates were Gaussian smoothed (std 20 ms) before trial-averaging. **b**. The Grid Evaluation Task. The participant attempted to move the cursor (white circle) to the cued target (green square) and click on it. **c**. Location of every click that was performed, relative to the current trial’s cued target (central black square), during blocks with the 6×6 grid (left) and with the 14×14 grid (right). Small gray squares indicate where the cursor began each trial, relative to the trial’s cued target. **d**. Timeline of the seventeen 3-minute Grid Evaluation Task blocks. Each point represents a trial and indicates the trial length and trial result (success or failure). Each gray region is a single block. **e**. T15’s online bitrate performance in the Grid Evaluation Task, compared to the highest-performing prior dPCG cursor control study. Circles are individual blocks (only shown for this study). Triangles are averages per participant (from this study and others).

Out of 1263 trials, T15 successfully navigated to and selected the correct target 1175 times (93%), selected the wrong target 88 times (7%), and reached the 10 second timeout without a selection 0 times (0%). There were also 6 clicks on the cued target while it was temporarily disabled, and 23 clicks not on any target. Of the 111 clicks not on the cued target, 107 (96%) of them were less than half a grid cell away from the cued target (Fig. 2c), suggesting that most errors were attributable to cursor decoding error as opposed to spontaneous click decodes.

### 3.3. Decoding contribution by brain area

T15’s microelectrode arrays (Fig. 3a and Supp. Fig. 2) were located in middle precentral gyrus (area 55b), ventral premotor cortex (area 6v), and primary motor cortex (area 4). In an offline analysis aimed at identifying which brain areas contributed to cursor decoding (Fig. 3b), we trained a decoder on each array alone as well as with each array excluded (i.e., only including the other three arrays). We trained each decoder using data from the Radial8 Calibration Task. We then replayed the neural data from the Grid Evaluation Task through each of these eight decoders and calculated each decoder’s average angular error per trial. Lower average angular error corresponds to better performance, but note that average angular error need not be near 0° to yield good cursor trajectories because the smoothing across time bins ameliorates many decoding errors. The decoder trained on d6v alone (40° angular error) performed best, almost matching the decoder trained on all arrays combined (39°). The decoder trained on 55b alone (88°) performed near chance-level, and excluding this array did not significantly impact offline decoding. Of note, 55b and v6v were the best arrays for speech decoding in Card et al. 2024. The decoders trained on the area 4 array alone (61°) and v6v alone (75°) had intermediate performance, but excluding either one did not significantly impact offline decoding.

**Figure 3.**
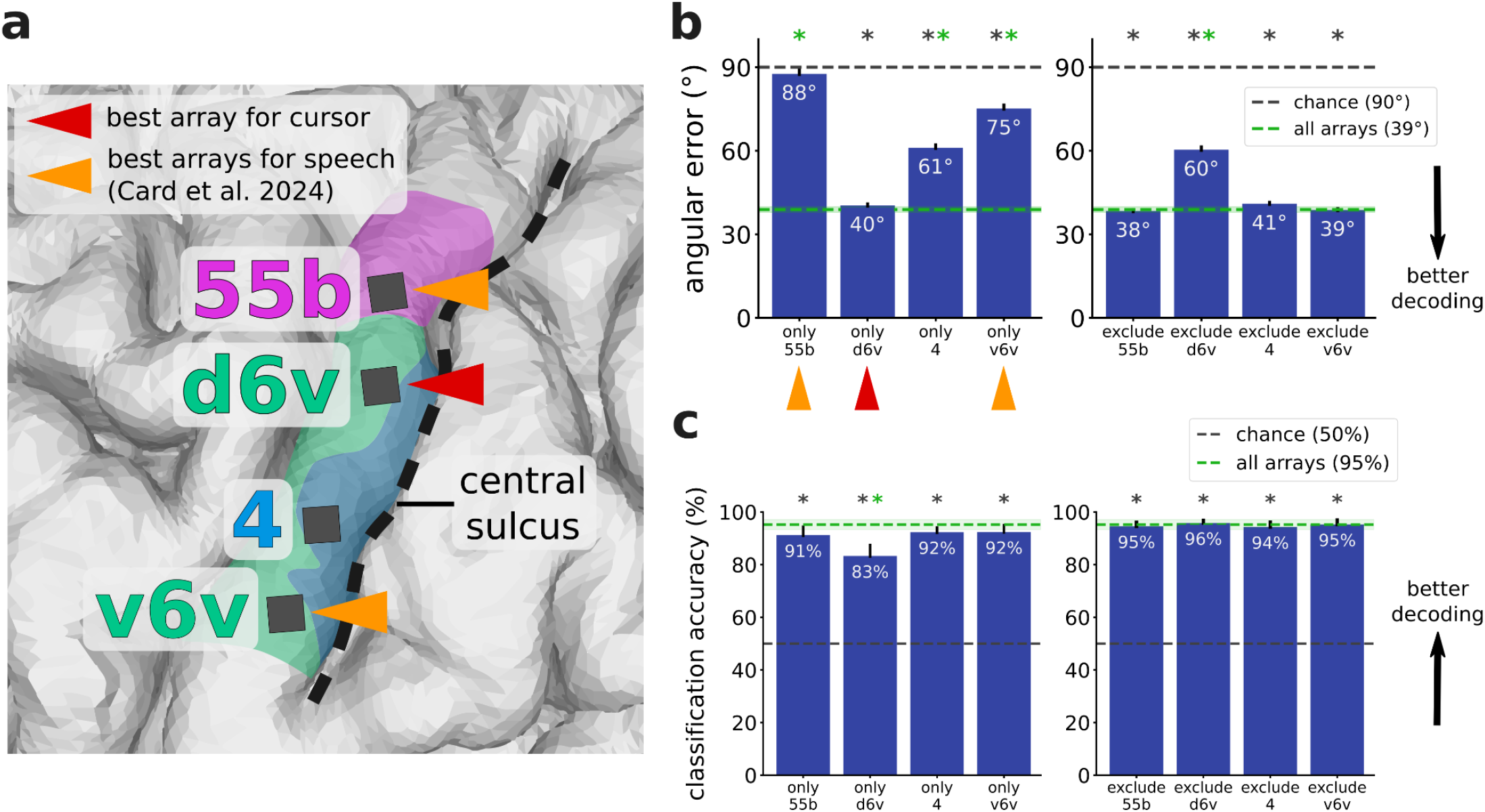
The dorsal 6v array contributed the most to cursor velocity decoding. **a**. Zoomed-in view of T15’s array locations shown in Fig. 1b. Triangles indicate arrays providing the best decoding performance for speech (orange) and for cursor control (crimson). The best speech arrays were identified in Card et al. 2024. **b**. Offline analysis of cursor decoders trained using neural features from each array alone (left) and with each array excluded (right), evaluated according to average angular error between the vector toward the target and the vector predicted by the decoder. The decoder trained on d6v alone performed almost the same as the decoder trained on all arrays. Error bars and the green shaded region indicate 95% CI. Stars indicate significant difference from chance performance (gray star, *p*<0.01, one-sample t-test, Bonferroni correction) or from performance using all arrays (green star, *p*<0.01, two-sample t-test, Bonferroni correction). **c**. Offline classification of click vs. non-click time windows using neural features from each array alone (left) and with each array excluded (right), evaluated with classification accuracy. Each of the four array locations contained click information. Error bars and the green shaded region indicate 95% CI. Stars indicate significant difference from chance performance (gray star, *p*<0.01, one-sample t-test, Bonferroni correction) or from performance using all arrays (green star, *p*<0.01, two-sample t-test, Bonferroni correction). Per-electrode contribution maps for cursor and click are shown in Supp. Fig. 2.

We ran an analogous analysis to identify which brain areas contained click-related information (Fig. 3c). In the case of cursor control, we had assumed the participant’s intention at each moment was to move toward the target. Here in the case of click, the participant’s intention (i.e., when exactly they attempt to click) was not known precisely because they may not have attempted to click the moment they reached the cued target; instead, they might have first tried to bring the cursor to a halt to avoid clicking the wrong target. So for this analysis, we assumed a high reliability of the click decoder used online during the Grid Evaluation Task and treated clicks decoded online as the true click intention (Fig. 2c). From every trial in the Grid Evaluation Task, we took the time window from 110-10 ms before the trial ended (which was always a click) and labeled it ‘click’, and took the time window from 800-700 ms before the trial ended and labeled it ‘non-click’. For each array alone and with each array excluded, we trained a logistic regression model to classify these 100 ms time windows of neural features as ‘click’ or ‘non-click’. All four of T15’s arrays yielded classification accuracies significantly above chance, suggesting that click information was present in all four brain areas.

### 3.4. Cursor control while simultaneously speaking

Because ventral precentral gyrus is typically associated with speech motor control, we investigated whether speaking interfered with cursor control. After training a cursor decoder using the standard Radial8 Calibration Task, we engaged T15 in a Simultaneous Speech and Cursor Task (Fig. 4a), which is the Radial8 Calibration Task with simultaneous speech cues. While the participant moved the cursor to each target, some trials (the ‘yes beep’ trials) contained an auditory beep (the ‘speech go cue’) at a random time to prompt the participant to speak a specific word, and other trials (the ‘no beep’ trials) contained no beep. To control for the added cognitive demands of attending to the beep instructions, in some blocks (the ‘never speak’ blocks) the participant was instructed not to speak at all even if they heard a beep, as opposed to the default behavior (the ‘speak if beep’ blocks) where the participant was supposed to speak once they heard the beep. We observed a variety of single-electrode tuning patterns: cursor movements only, speaking only, and both cursor and speech (Fig. 4b).

**Figure 4.**
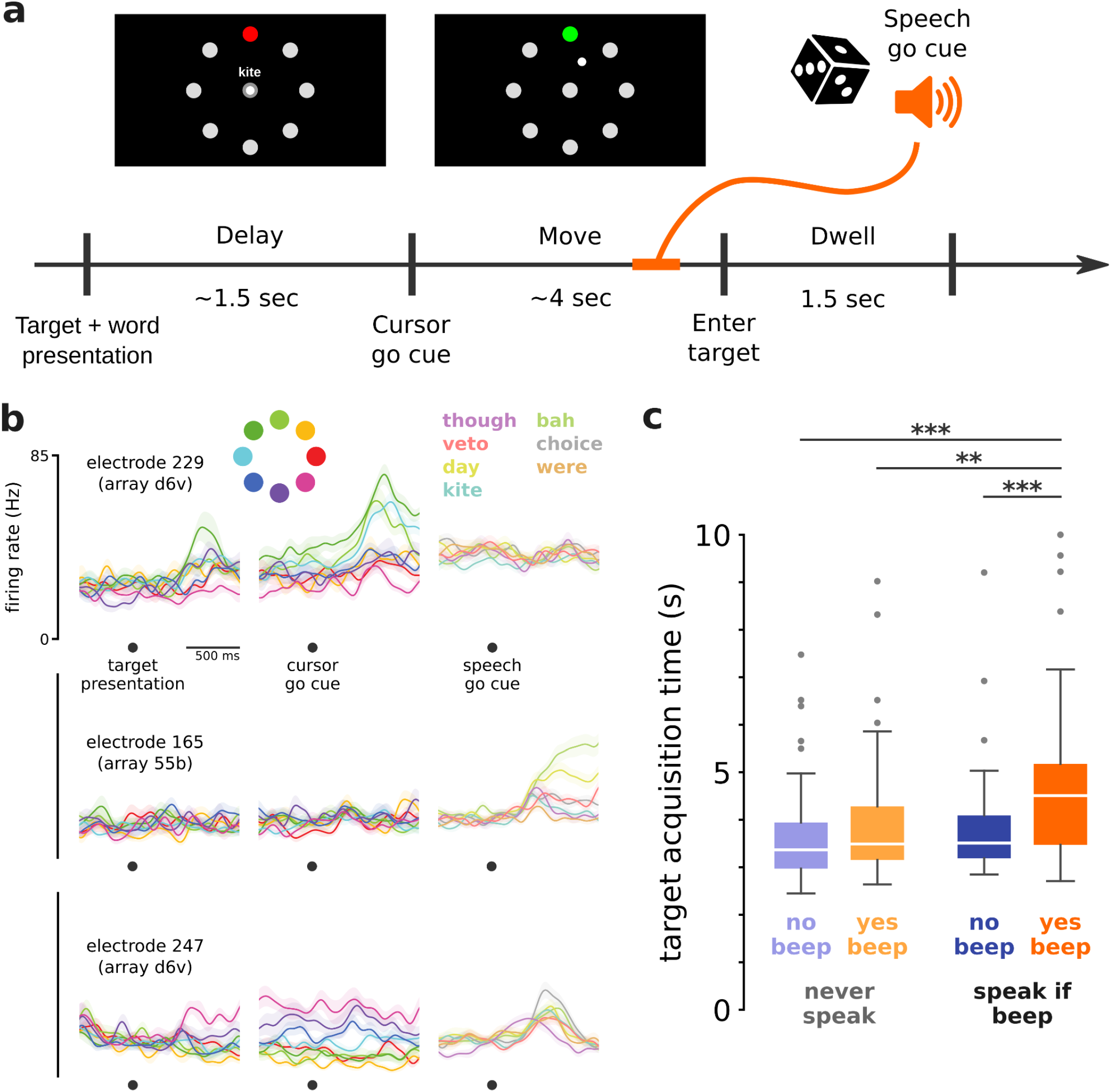
Cursor control was impacted by simultaneously speaking. **a**. Overview of the Simultaneous Speech and Cursor Task. Each trial may or may not have a speech go cue (an auditory beep), which could occur during the move period or dwell period. **b**. Trial-averaged firing rates (mean ± s.e.) recorded from three example electrodes during the Simultaneous Speech and Cursor Task. Activity during cursor control alone (left and center columns) is aligned first to target presentation and then to cursor go cue, and is colored by cued target direction. Activity during simultaneous cursor control and attempted speech (right column) is aligned to speech go cue, and is colored by prompted word. Firing rates were Gaussian smoothed (std 50 ms) before trial-averaging. **c**. Comparison of target acquisition times in different trial conditions in the Simultaneous Speech and Cursor Task. Target acquisition times in the only condition requiring speaking were significantly longer than the other conditions (* *p*<0.05, ** *p*<0.01, *** *p*<0.001, rank-sum test, Bonferroni correction).

For each of the four conditions (each combination of the two trial types and two block types introduced above), we measured T15’s target acquisition times (Fig. 4c). Target acquisition times for the only condition with speech (‘speak if beep - yes beep’) were significantly longer (median 4.51 seconds) than in each of the conditions without speech: ‘speak if beep - no beep’ (3.51 seconds), ‘never speak - yes beep’ (3.49 seconds), and ‘never speak - no beep’ (3.37 seconds). This suggests that speech production did interfere with vPCG cursor control, in this scenario where speech was not specifically accounted for in the decoder or training process. Though many electrodes showed modulation to the task’s auditory stimuli, trials with the speech go cue but no speech (‘never speak - yes beep’) did not have significantly longer target acquisition times, suggesting that it was indeed the production of speech (and not the perceived beep and auditory feedback) which interfered with cursor control.

### 3.5. Personal use of desktop computer

In a prior study, T15 used the BCI system’s speech decoding capabilities in a personal use setting.^22^ In the current study, we added the ability to neurally control T15’s personal computer’s (Mac) mouse cursor (Fig. 5). Video of T15 using the cursor BCI to browse Netflix and play the New York Times Spelling Bee are shown in Supp. Movie 5 and Supp. Movie 6. Without a cursor BCI, T15 controlled his computer with a gyroscopic head mouse (Zono 2, Quha), or with the aid of a care partner or family member.

**Figure 5.**
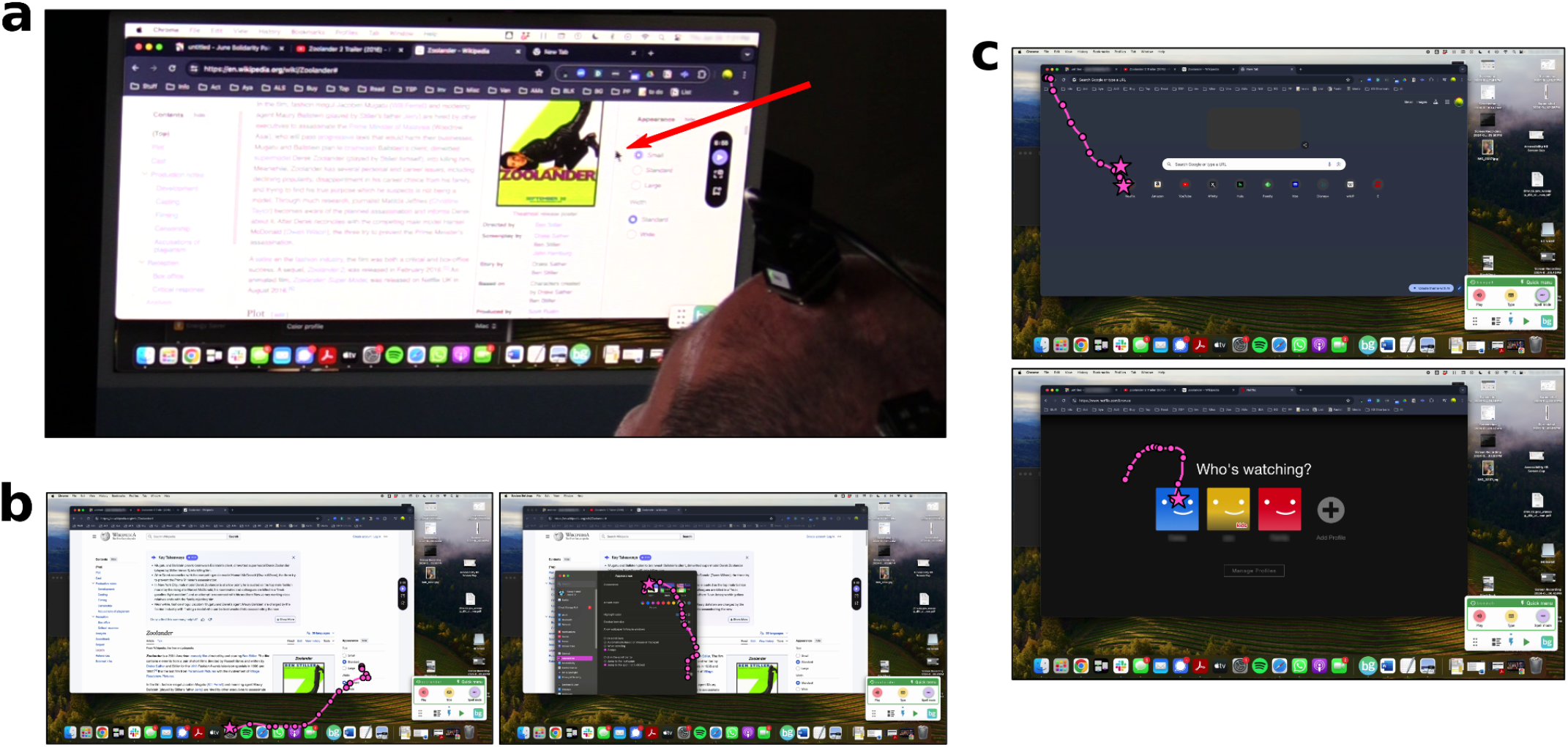
Participant T15 controlled his personal desktop computer with the cursor BCI. **a**.Over-the-shoulder view of T15 neurally controlling the mouse cursor on his personal computer. The red arrow points to the cursor. **b-c**. Screenshots of T15’s personal computer usage, with cursor trajectories (pink lines) overlaid. Cursor position every 250 ms (circles) and clicks (stars) are also drawn. In b., T15 first opened the Settings application (left) and then switched his computer to Light Mode (right). In c., T15 opened Netflix from Chrome’s New Tab menu (top) and then selected his Netflix user (bottom).

## 4. Discussion

We demonstrated a rapidly-calibrating, accurate cursor BCI driven by vPCG, an area canonically associated with speech and orofacial movements. The finding that the cursor BCI could be controlled with non-speech, non-orofacial imagery aligns well with accumulating evidence that the whole body is represented in a distributed fashion along the precentral gyrus.^24–27^ Furthermore, this study exemplifies how to leverage this modern view of motor cortex to build multi-modal implanted BCIs using fewer arrays and implant sites. We note, however, that cursor velocity decoding relied heavily on a single array (in the dorsal aspect of area 6v). Achieving similar cursor performance from vPCG may require specifically seeking out this area preoperatively, and it is not yet known whether a similar motor map is present (and can be reliably located) in other BCI users, as this study includes a single participant.

### 4.1. Cursor decoding performance

After optimization, the cursor BCI achieved an average bitrate of 2.90 bps. Compared to the highest-performing previously published dPCG cursor BCI,^10^ this is higher than participant T6 (2.2 bps) and T7 (1.4 bps) but lower than T5 (4.16 bps). It is also lower than unpublished results from participant P1 in Neuralink’s PRIME clinical trial.^43^ From a pragmatic perspective, the cursor BCI yielded ample performance for a participant to control their own computer for their own purposes, and the participant used the BCI both for work and for leisure.

There also may be opportunities to improve performance. Here we used linear decoders that assume static neural tuning properties because these techniques were simple and sufficient. However, a growing body of evidence for the factor-dynamics perspective^44^ (which emphasizes the heterogeneous, dynamic, and complex tuning properties present at the single-unit level) suggests that more expressive decoding architectures such as recurrent neural networks (RNNs)^16,45,46^ may yield higher cursor decoding performance.

### 4.2. Motor imageries driving cursor control

The participant successfully controlled the cursor under a variety of instructed cursor imageries covering different body parts (hand, tongue, head). However, for some imageries and sessions the actual movements attempted by the participant may not have matched the instruction, for two reasons.

First, in early sessions the participant visibly moved his head along with the cursor due to familiarity with using a head mouse. We asked the participant if he could keep his head still but he found it too uncomfortable to suppress head movement while trying to control the cursor. We revisited the head movement topic in a later session, and by relaxing his head on his head rest the participant was able to comfortably keep his head mostly stationary (he still made intermittent, small head movements to view the cursor and target). In this regime, cursor control continued to work, so we believe cursor control did not rely on overt head movements.

Second, the participant frequently described his method of moving the cursor as “intuition”, even for blocks where there was a specific body part instructed. In his own words, “I have to use The Force” (referring to the telekinetic ability from Star Wars films). Such effector-agnostic, subjective experiences during neuroprosthesis control are widespread in past BCI studies (although not formally reported), and present a challenge for interpreting results that rely on the instructed imagery. These caveats preempt claims comparing specific motor imageries, but do not detract from the practicality of the cursor BCI demonstrated. Future work could more carefully investigate representations of different body parts in this brain area.

### 4.3. Interference from speech-related neural activity

Speech interfered with cursor control, in contrast to when cursor control is driven by dPCG.^28^ For now, this may encourage a BCI user to attempt to speak and control the cursor sequentially instead of in parallel. This did not impede the participant from using the multi-modal BCI for daily life activities in which he moved the cursor of his personal computer and used the speech BCI for text entry. Of note, most able-bodied computer users use a mouse and keyboard sequentially. Nevertheless, cursor control that is robust to simultaneous speech could improve the user experience. This may be possible to achieve with modifications only to the calibration protocol, without needing to modify the decoder architecture, e.g., by prompting the user to speak at random times during the cursor calibration task so that the cursor decoder learns to filter out speech-related neural modulation. Another strategy could be to modify the cursor decoder to explicitly account for speech-related neural modulation, using data either from a modified cursor calibration task or from speech-only windows collected previously during speech BCI usage.

## 5. Conclusion

As speech BCIs driven by vPCG mature and proliferate, it is valuable to identify what other BCI applications vPCG can enable. Here, a participant with ALS who previously achieved highly accurate speech decoding^22^ was able to control his own computer as if using a computer mouse. This marks an important step toward delivering high-performing, multi-modal BCIs to people with paralysis.

## Acknowledgements

We thank participant T15, his family, and his carepartners for their dedicated contributions to this research.

This work was supported by the National Science Foundation Research Traineeship NRT program Award #2152260, and the ARCS Foundation (Singer-Clark); the Office of the Assistant Secretary of Defense for Health Affairs through the Amyotrophic Lateral Sclerosis Research Program under award number AL220043; DP2 from the NIH Office of the Director and managed by NIDCD (1DP2DC021055); Searle Scholars Program; and a Career Award at the Scientific Interface from the Burroughs Wellcome Fund (Stavisky); The content is solely the responsibility of the authors and does not necessarily represent the official views of the National Institutes of Health, or the Department of Veterans Affairs, or the United States Government.

## Supplementary Materials

**Supplementary Movie 1. Radial8 Calibration Task**. Participant T15 performing the Radial8 Calibration Task. Cursor movement and click were in full closed-loop neural control. This was the third of three consecutive calibration blocks, so the cursor decoder and click decoder being used were trained on the previous two calibration blocks and continually updated throughout the video. There was no specific instructed imagery (i.e., T15 described his method of control as “intuition” for both cursor and click).

Link to view online: https://ucdavis.box.com/s/aywxbzq4l73f23e58kzwvtpa0hd698iw

**Supplementary Movie 2. Grid Evaluation Task**. Participant T15 performing the Grid Evaluation Task. Cursor movement and click were in full closed-loop neural control. There was no specific instructed imagery (i.e., T15 described his method of control as “intuition” for both cursor and click). Link to view online: https://ucdavis.box.com/s/i5z9oznksdea6s46afxgw5fxjuikl1hd

**Supplementary Movie 3. Simultaneous Speech and Cursor Task**. Participant T15 performing the Simultaneous Speech and Cursor Task. Cursor movement was in full closed-loop neural control. The block type was ‘speak if beep’, so T15 attempted to speak the words (inaudible in this video) when he heard the beep (the ‘speech go cue’). The instructed cursor imagery was ‘Right Hand Mouse’. Targets were selected by dwelling the cursor on them (i.e., there was no clicking).

Link to view online: https://ucdavis.box.com/s/jksikp6t3rhi70r87678dun70zw29q2v

**Supplementary Movie 4. Rapid calibration during first-ever cursor BCI usage**. Participant T15’s first 3 minutes of ever using a cursor BCI. As described in Section 2.3.1, the task began in open-loop and gradually transitioned to closed-loop as a decoder being trained in the background improved. From the 35 second mark on, T15 had full closed-loop neural control of the cursor. The instructed cursor imagery was ‘Right Hand Mouse’, but T15 also moved his head along with the cursor during this block. T15 had familiarity with using a head mouse from past experience.

Link to view online: https://ucdavis.box.com/s/uka4ddljzqmhhphihsim0qr9gezudmqi

**Supplementary Movie 5. Personal use (opening and browsing Netflix)**. Over-the-shoulder view of participant T15 using the cursor BCI to control his personal computer’s mouse cursor, with screen capture of his monitor overlaid. In this clip, T15 opened Netflix in a new browser tab, selected his own user account, scrolled down the page by clicking the scroll bar, and started watching a TV show.

Link to view online: https://ucdavis.box.com/s/x39nfzujcfdf0czhj3h43r720rgk4lts

**Supplementary Movie 6. Personal use (playing the New York Times Spelling Bee)**.

Over-the-shoulder view of participant T15 using the cursor BCI to control his personal computer’s mouse cursor, with screen capture of his monitor overlaid. In this clip, T15 typed and submitted words in the New York Times Spelling Bee web game by clicking each letter and clicking the word submission button.

Link to view online: https://ucdavis.box.com/s/ac6pymrvvenydg5smhbn6ipfguojkuzu

**Supplementary Figure 1.**
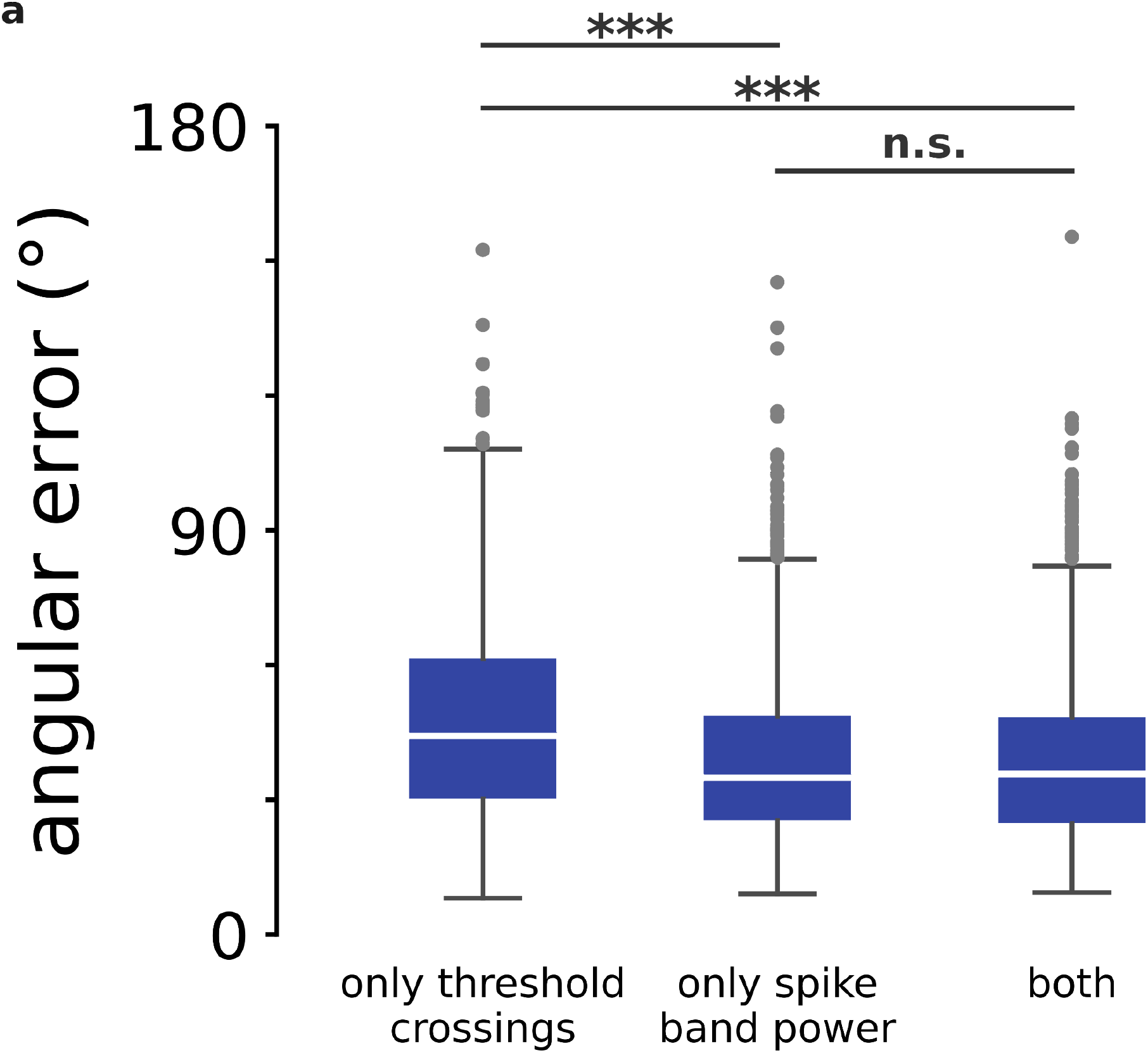
Comparison of neural feature types for cursor decoding. **a**. Angular error per trial computed offline under three different choices of neural features used for decoding: threshold crossings, spike band power, and both combined. Input vectors to the training and decoding algorithms were 512-D when using both types of neural features (as was done online in this study) and 256-D when using each feature type alone. There was no significant difference between using spike band power alone vs. using spike band power and threshold crossings together (* *p*<0.05, ** *p*<0.01, *** *p*<0.001, rank-sum test, Bonferroni correction).

**Supplementary Figure 2.**
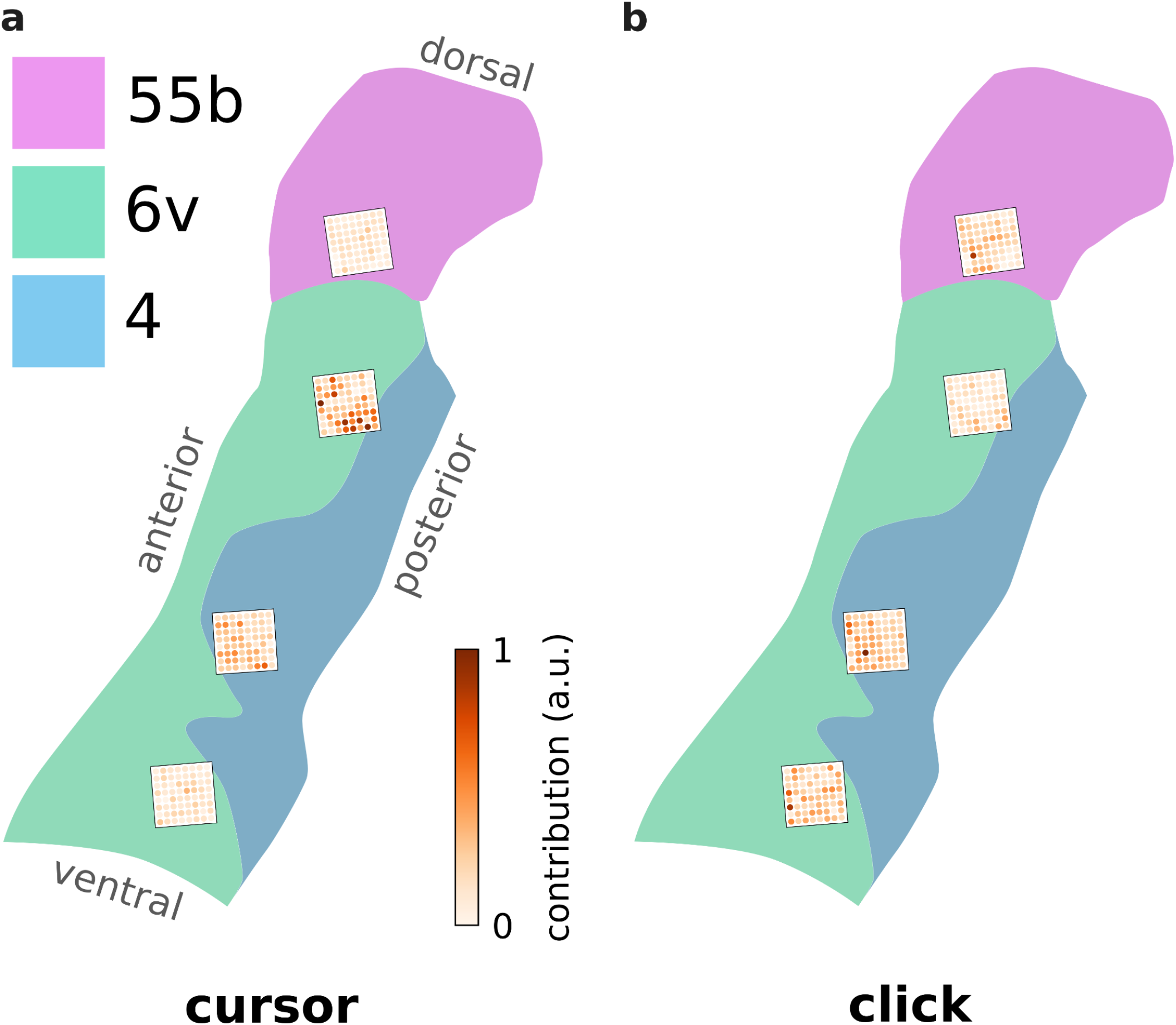
Microelectrode contributions to cursor and click. **a**. Individual microelectrodes, colored by their relative contribution to the cursor decoder’s output vectors during the Grid Evaluation Task. The arrays are overlaid on the HCP parcellation results for T15’s vPCG. **b**. Same as panel a, but for click decoding. Electrodes are colored by the portion they contributed to the click decoder’s class probability estimates. See Sections 2.4.2 and 2.5.1 for how each electrode’s decoder contributions were calculated.

## References

1. Serruya, M. D., Hatsopoulos, N. G., Paninski, L., Fellows, M. R. & Donoghue, J. P. Instant neural control of a movement signal. Nature 416, 141–142 (2002).

2. Carmena, J. M. et al. Learning to Control a Brain–Machine Interface for Reaching and Grasping by Primates. PLOS Biol. 1, e42 (2003).

3. Nuyujukian, P., Fan, J. M., Kao, J. C., Ryu, S. I. & Shenoy, K. V. A High-Performance Keyboard Neural Prosthesis Enabled by Task Optimization. IEEE Trans. Biomed. Eng. 62, 21–29 (2015).

4. Leuthardt, E. C., Schalk, G., Wolpaw, J. R., Ojemann, J. G. & Moran, D. W. A brain–computer interface using electrocorticographic signals in humans. J. Neural Eng. 1, 63–71 (2004).

5. Hochberg, L. R. et al. Neuronal ensemble control of prosthetic devices by a human with tetraplegia. Nature 442, 164–171 (2006).

6. Kim, S.-P. et al. Point-and-Click Cursor Control With an Intracortical Neural Interface System by Humans With Tetraplegia. IEEE Trans. Neural Syst. Rehabil. Eng. 19, 193–203 (2011).

7. Orsborn, A. L. et al. Closed-Loop Decoder Adaptation Shapes Neural Plasticity for Skillful Neuroprosthetic Control. Neuron 82, 1380–1393 (2014).

8. Jarosiewicz, B. et al. Virtual typing by people with tetraplegia using a self-calibrating intracortical brain-computer interface. Sci. Transl. Med. 7, 313ra179–313ra179 (2015).

9. Vansteensel, M. J. et al. Fully Implanted Brain–Computer Interface in a Locked-In Patient with ALS. N. Engl. J. Med. 375, 2060–2066 (2016).

10. Pandarinath, C. et al. High performance communication by people with paralysis using an intracortical brain-computer interface. eLife 6, e18554 (2017).

11. Brandman, D. M. et al. Rapid calibration of an intracortical brain–computer interface for people with tetraplegia. J. Neural Eng. 15, 026007 (2018).

12. Nuyujukian, P. et al. Cortical control of a tablet computer by people with paralysis. PLOS ONE 13, e0204566 (2018).

13. Willett, F. R. et al. Principled BCI Decoder Design and Parameter Selection Using a Feedback Control Model. Sci. Rep. 9, 8881 (2019).

14. Weiss, J. M., Gaunt, R. A., Franklin, R., Boninger, M. L. & Collinger, J. L. Demonstration of a portable intracortical brain-computer interface. Brain-Comput. Interfaces 6, 106–117 (2019).

15. Dekleva, B. M., Weiss, J. M., Boninger, M. L. & Collinger, J. L. Generalizable cursor click decoding using grasp-related neural transients. J. Neural Eng. 18, 0460e9 (2021).

16. Deo, D. R. et al. Brain control of bimanual movement enabled by recurrent neural networks. Sci. Rep. 14, 1598 (2024).

17. Candrea, D. N. et al. A click-based electrocorticographic brain-computer interface enables long-term high-performance switch scan spelling. Commun. Med. 4, 1–14 (2024).

18. Moses, D. A. et al. Neuroprosthesis for Decoding Speech in a Paralyzed Person with Anarthria. N. Engl. J. Med. 385, 217–227 (2021).

19. Luo, S. et al. Stable Decoding from a Speech BCI Enables Control for an Individual with ALS without Recalibration for 3 Months. Adv. Sci. 10, 2304853 (2023).

20. Metzger, S. L. et al. A high-performance neuroprosthesis for speech decoding and avatar control. Nature 620, 1037–1046 (2023).

21. Willett, F. R. et al. A high-performance speech neuroprosthesis. Nature 620, 1031–1036 (2023).

22. Card, N. S. et al. An Accurate and Rapidly Calibrating Speech Neuroprosthesis. N. Engl. J. Med. 391, 609–618 (2024).

23. Wairagkar, M. et al. An instantaneous voice synthesis neuroprosthesis. 2024.08.14.607690 Preprint at 10.1101/2024.08.14.607690 (2024).

24. Stavisky, S. D. et al. Neural ensemble dynamics in dorsal motor cortex during speech in people with paralysis. eLife 8, e46015 (2019).

25. Wilson, G. H. et al. Decoding spoken English from intracortical electrode arrays in dorsal precentral gyrus. J. Neural Eng. 17, 066007 (2020).

26. Willett, F. R. et al. Hand Knob Area of Premotor Cortex Represents the Whole Body in a Compositional Way. Cell 181, 396-409.e26 (2020).

27. Deo, D. R. et al. A mosaic of whole-body representations in human motor cortex. 2024.09.14.613041 Preprint at 10.1101/2024.09.14.613041 (2024).

28. Stavisky, S. D. et al. Speech-related dorsal motor cortex activity does not interfere with iBCI cursor control. J. Neural Eng. 17, 016049 (2020).

29. Glasser, M. F. et al. A multi-modal parcellation of human cerebral cortex. Nature 536, 171–178 (2016).

30. Young, D. et al. Signal processing methods for reducing artifacts in microelectrode brain recordings caused by functional electrical stimulation. J. Neural Eng. 15, 026014 (2018).

31. Simeral, J. D., Kim, S.-P., Black, M. J., Donoghue, J. P. & Hochberg, L. R. Neural control of cursor trajectory and click by a human with tetraplegia 1000 days after implant of an intracortical microelectrode array. J. Neural Eng. 8, 025027 (2011).

32. Orsborn, A. L., Dangi, S., Moorman, H. G. & Carmena, J. M. Closed-Loop Decoder Adaptation on Intermediate Time-Scales Facilitates Rapid BMI Performance Improvements Independent of Decoder Initialization Conditions. IEEE Trans. Neural Syst. Rehabil. Eng. 20, 468–477 (2012).

33. Kao, J. C., Nuyujukian, P., Ryu, S. I. & Shenoy, K. V. A High-Performance Neural Prosthesis Incorporating Discrete State Selection With Hidden Markov Models. IEEE Trans. Biomed. Eng. 64, 935–945 (2017).

34. Nuyujukian, P. et al. Performance sustaining intracortical neural prostheses. J. Neural Eng. 11, 066003 (2014).

35. Willett, F. R. et al. Feedback control policies employed by people using intracortical brain–computer interfaces. J. Neural Eng. 14, 016001 (2016).

36. Georgopoulos, A., Kalaska, J., Caminiti, R. & Massey, J. On the relations between the direction of two-dimensional arm movements and cell discharge in primate motor cortex. J. Neurosci. 2, 1527–1537 (1982).

37. Willett, F. R. et al. A Comparison of Intention Estimation Methods for Decoder Calibration in Intracortical Brain–Computer Interfaces. IEEE Trans. Biomed. Eng. 65, 2066–2078 (2018).

38. Gilja, V. et al. A high-performance neural prosthesis enabled by control algorithm design. Nat. Neurosci. 15, 1752–1757 (2012).

39. Jarosiewicz, B. et al. Advantages of closed-loop calibration in intracortical brain–computer interfaces for people with tetraplegia. J. Neural Eng. 10, 046012 (2013).

40. Fan, J. M. et al. Intention estimation in brain–machine interfaces. J. Neural Eng. 11, 016004 (2014).

41. Ali, Y. H. et al. BRAND: a platform for closed-loop experiments with deep network models. J. Neural Eng. 21, 026046 (2024).

42. Pedregosa, F. et al. Scikit-learn: Machine Learning in Python. Mach. Learn. PYTHON.

43. Neuralink. PRIME Study Progress Update — User Experience. Neuralink Blog https://neuralink.com/blog/prime-study-progress-update-user-experience/ (2024).

44. Churchland, M. M. & Shenoy, K. V. Preparatory activity and the expansive null-space. Nat. Rev. Neurosci. 1–24 (2024) doi:10.1038/s41583-024-00796-z.

45. Sussillo, D., Stavisky, S. D., Kao, J. C., Ryu, S. I. & Shenoy, K. V. Making brain–machine interfaces robust to future neural variability. Nat. Commun. 7, 13749 (2016).

46. Hosman, T., Pun, T. K., Kapitonava, A., Simeral, J. D. & Hochberg, L. R. Months-long High-performance Fixed LSTM Decoder for Cursor Control in Human Intracortical Brain-computer Interfaces. in 2023 11th International IEEE/EMBS Conference on Neural Engineering (NER) 1–5 (2023). doi:10.1109/NER52421.2023.10123740.

